# Using lightsheet microscopy to investigate the initial lymphatic network in the murine knee joints

**DOI:** 10.1101/2025.03.21.644620

**Authors:** L. Xing, X. Lin

**Affiliations:** Dept of Pathology and Laboratory Medicine, University of Rochester Medical Center, USA; Orthopaedics and Rehabilitation, Center for Musculoskeletal Research, University of Rochester Medical Center, USA; Biostatistics and Computational Biology, University of Rochester Medical Center, USA

## Abstract

Introduction. Lymphatic vasculature in mouse joints is difficult to study due to its vast anatomical variation. We reported increased lymphatic capillaries in post-traumatic OA (PTOA) murine joint based on 2D immunostaining. However, a gap of knowledge with this approach is that histology highly subjected to histological sampling, which might not be appropriate for a highly variable system such as the lymphatic vasculature. We hypothesize that using lightsheet microscopy would reveal the detailed structure of the initial lymphatic network.

Methods. We established lightsheet imaging protocol and four quantifiable outcome parameters for visualizing and analyzing the initial lymphatic network in young C57/B6 wild type mouse knees. Lymphatic vessels were identified by staining samples with lymphatic endothelial marker LYVE1. We validated our lightsheet imaging protocol in mouse knee joints following PTOA surgery and knee joints received sham operation.

Results. We used the histological landmarks of growth plates and menisci shape for consistent data collection for the 3D reconstruction of initial lymphatic network. Comparing to sham joints, PTOA joints have less mean branches length (58.91± 6.50 vs. 146.29 ± 2.48 μm in sham, *p*<0.0001), and mean branch diameter (12.25± 0.80] vs. 28.51 ± 4.99 μm in sham, *p*=0.0051), but higher total branching points (378± 239 vs. 69 ± 17 μm in sham, *p*=0.020), and total initial lymphatic network volume (51247± 14239 vs. 1019289 ± 544458 μm^3^ in sham, *p*=0.037). Conclusion. Lightsheet imaging of murine knee joints is a powerful and promising tool to study the joint lymphatic system and arthritis. Similar to previous report, the PTOA joint has increased branching with smaller and shorter branches.

## Introduction

The lymphatic circulation was enigmatic due to its anatomical heterogeneity and lack of lymphatic vessel specific markers, until the discovery of Podoplanin (PDPN)[1], Lymphatic vessel endothelial hyaluronan receptor 1 (LYVE1)[2], and Prospero homeobox protein 1 (PROX1)[3] as specific markers of lymphatic endothelial cells (LECs) were discovered around the 2000s, allowing the discovery of the lymphatic system in various tissues, including the heart, eye, brain, and joints. The initial lymphatic capillary network in the tissues merge into collecting lymphatic vessels that connect the lymph nodes and drain into the vein, completing the lymphatic circulation[4, 5]. The lymphatic circulation is critical for maintaining fluid homeostasis.

The synovial lymphatic system (SLS) is discovered and characterized by combining innovative imaging modalities and LV markers: 1) Unlike the blood circulation where the heart is a central pump to propel the blood flow, the lymphatic flow relies on alternate contraction of the LMC and LV valves to prevent retrograde flow. When injecting a small molecule indocyanine green (ICG, molecular weight: 751 Da) to the synovial tissue, ICG enters both the blood and lymphatic circulation but can be used for in vivo imaging of the SLS due to the slower flow in the lymphatics compared to the blood circulation. Thus, we developed the near-infrared and ICG (NIR-ICG) lymphatic imaging[6-11] to monitor lymphatic vessel (LV) contraction and clearance from joints to draining lymph nodes. 2) Based on the cellular composition of the capillary and collecting LVs, which is LEC or LEC+LMC respectively, we developed immunofluorescence staining/whole slide imaging (IF/WSI) to identify and qualify lymphatic vessels in mouse joints[12, 13]. Capillary LV is detected by LEC specific markers including LYVE1 or PROX1. Collecting LV is detected by LYVE1/PROX1+αSMA double staining. 3) Unlike the blood capillaries, capillary LVs are highly fenestrated and non-selectively allows the exchange of macromolecules. Utilizing this feature, 70kDa fluorophore conjugated Dextran was used for angiography and lymphatic imaging[14], when injected intravenously or to the tissue, respectively. We developed IVIS-Dextran imaging for the SLS, and established three parameters [15], the SLS clearance, the influx of the capillary LVs, the capacity of DLNs. The SLS is associated with the physiological and pathological conditions such as rheumatoid arthritis, post traumatic osteoarthritis, natural aging and age-related osteoarthritis. The initial lymphatic network of the SLS is preset in the synovium and surrounding soft tissues around the joint capsule, ligaments, fat pads, and muscles of normal knees. Initial lymphatic network merge into collecting LVs and drains to the DLNs, which are iliac LNs for the knee and popliteal LNs for the ankle[9, 16]. The SLS removes catabolic molecules, including metalloproteinases and inflammatory cytokines, from the joint to maintain homeostasis and protects against arthritis progression[8, 15]. Damage to the SLS exacerbates arthritis and, vice versa, enhancement of the SLS attenuates arthritis[7, 10, 15, 17].

While we have gained significant understanding of the SLS, a gap in knowledge of the link between anatomic distribution/histomorphometry of the SLS LVs and SLS function limits comprehensive understanding of the role of SLS in the joint. NIR-ICG and IVIS-Dextran in vivo imaging do not have the sufficient resolution to distinguish individual vessels in the joint, and can only assess functional changes. The IF/WSI imaging is limited by the area of interest due to photobleaching, and is highly influenced by histologic sampling of sectioned planes. Light sheet microscopy is a novel imaging technique that allows observation of the full depth of the joint tissue, compared to a maximum of 50 μm in traditional fluorescence imaging, and eliminate photobleaching as a limiting factor for the range of region of interest.

In this study, we establish the lightsheet imaging protocol and outcome measurements for the initial lymphatic network of murine knee joint, and validate our protocol using the post traumatic OA model, which has been reported to decreased SLS function and increased number of non-functional lymphatic capillaries.

## Materials and Methods

### Animals

5-month-old C57BL/6J male mice from National Cancer Institute Mouse Repository were used for destabilization of medial meniscus (DMM) or sham surgery procedure. Knee joints were harvested two weeks after the procedure. Mice were housed in two-way micro-isolator technique rodent rooms. All animal procedures were approved by the University Committee on Animal Research at the University of Rochester.

### Mouse knee tissue preparation and LYVE-1 immunostaining

Knee joints were dissected with femur and tibia. Muscles were thoroughly removed from the bone without disrupting the synovial tissues. Knee joints were fixed in 10% Formalin for 48 hours, decalcified in 14% EDTA solution for 10 days, and dehydrated. To label the initial lymphatic network with LYVE-1 antibodies, tissues were first underwent to 1) bleach (PBS 1hrx2, 50% methanol (in PBS) 1hr, 100% methanol 1hrx2, and 5% H2O2 in 20% DMSO/methanol (1 vol 30% H2O2/1 vol DMSO/4vol methanol, ice cold) at 4C overnight; 2) pretreatment (methanol 1hrx2, 20%DMSO/methanol 1hrx2, 80% methanol 1hr, 50% methanol 1hr, PBS overnight, PBS 1hr, PBS/02% Triton X-100 1hr, and PBS/0.2% Triton X-100/20%DMSO/0.3M glycine at 37C overnight); and 3) blockage (PBS/0.2% TritonX-100/10%DMSO/6% Donkey Serum at 37C overnight, PBS/0.2% Tween-20 with 10ug/mL heparin (PTwH) 1hr x2). Tissues were then stained with rabbit anti-mouse LYVE-1 Ab (1:?; Abcam, ab14917) in PTwH/5% DMSO/3% Donkey serum at 37C for 3 days, washed in PTwH for 1d (>10 washes with each>1hr), incubated in AlexaFluor 647 conjugated goat anti-Rabbit Ab (Invitrogen, A-21245) in PTwH/3% Donkey serum at 37C for 1 day, washed in PTwH for (>10 washes with each>1hr). Tissues were cleared in a simplified version of 3D imaging of solvent-cleared organs (3DISCO). Tissues were incubated in 10 ml of 50% v/v tetrahydrofuran/H2O (THF) (Sigma 186562-12×100ML) overnight, 1hr in 10 ml of 80% THF/H2O and twice 1hr in 100% THF, dichloromethane (DCM) (Sigma 270997-12×100ML) until they sink to the bottom of the vial, in 18 ml of dibenzyl ether (DBE) (Sigma 108014-1KG) until clear (2hr) and then stored in DBE at room temperature.

### Lightsheet imaging of the initial lymphatic network in the mouse knee joint

The cleared samples were imaged on the Bruker MuVi SPIM Microscope with the LuxControl software. The microscope was equipped with the Single Detection 4x CS octagon with Nikon Plan Fluor 4x/0.2 NA illumination objectives and an Olympus 4x/0.28NA detection objective. The microscope was fitted with the 405nm, 445nm, 488nm, 515nm, 561nm and 642nm diode laser lines. The images were captured with the Hamamatsu Orca Flash 4.0 V3 sCMOS camera with a final image resolution of ?nm in the x and y dimensions. For image acquisition, cleared samples were mounted onto the sample holder adaptor and were gently immersed in DBE in the imaging chamber. Samples were excited with a scanned light sheet (beam expander 2.8um) of 488nm to detect autofluorescence and 642nm to detect LYVE1 signal by immunostaining. The raw data were processed by Luxendo Processor and then converted by Imaris File converter (version10.2.0, Oxford Instrument) and then analyzed by Imaris software (version 10.2.0, Oxford Instrument).

### Imaging Analysis

Slices/Z-stacks of images acquired on the light sheet were processed and reconstructed in three dimensions in the 488nm and 642nm channel with the Imaris software (version 10.2.0). To generate consistent images and analysis, all samples were thresholded with with an intensity range of “Min” = 500 to “Max” = 2000. For branching analysis, a mask excluding any muscle tissue was created. Using the “Autopath” algorithm in the “Filament” module, where a machine-learning based algorithm produced a filament model with loops based on local intensity contrast in the 642nm channel. The minimum and maximum diameter of a “filament” were defined by the user based on the smallest and largest visible vessels seen on series of 2D images. Notably, the sham joints have a diameter range of 24.73 ± 4.45 – 45.63 ± 2.29 μm, while the PTOA joints have a significantly smaller diameter range of 6.40 ± 0.75 – 43.60 ± 3.99 μm (Supplementary Fig. 1). The algorithm then predict filaments in a user-defined ROI as the training set, which allows the user to optimize the filament model by discarding any predicted filaments that did not overlap with LYVE1+ staining. The algorithm then proceeded to process the whole image based on the training set. Samples were numbered randomly before quantification, the number of branch points, the mean branch length and diameter, and the total volume were acquired using the “Statistics” function.

### Statistical analysis

Sample sizes were determined by a priori power calculations using the G*power software (version 3.1). For assessing the difference of initial lymphatic network between the sham and DMM mice, the sample size n>=3 was calculated given expected means capillary LV area 25 (5) mm^2^ in sham knee and 75 (20) mm^2^ in PTOA knee [13], with an alpha level of 0.05 and a power of 80%, for a two-sample t-test with unequal variance. Was performed using GraphPad Prism 5 software (GraphPad Software Inc., San Diego, CA, USA). Statistical analysis was performed using GraphPad Prism 10 software (GraphPad Software Inc., San Diego, CA, USA). Data are presented as mean ± SD. Comparisons between 2 groups were analyzed using a 2-tailed unpaired Student’s t-test. For non parametric data, comparisons between 2 groups were analyzed using the Mann Whitney U test.

## Results

### Using tissue autofluorescence to establish the region of interest for the synovial lymphatic lightsheet microscopy

We established a working protocol to visualize initial lymphatic vasculature in mouse knee joints using anti-LYVE-1 antibody (Fig.1). Since lymphatic vessels are composed lymphatic capillaries and mature (collecting lymphatic vessels) which cannot be distinguish by LYVE-1 staining, we named all LYVE-1+ vessels as initial lymphatic vessels in the current study. We mounted the joint tissue onto an open-cassette sample holder with the sagittal plane of the joint parallel to the lightsheets and the medial side of the joint entering the lightsheet first, similar to the way we perform histology sectioning of the joint. We collected data in the 488nm channel, where we utilized the tissue autofluorescence to visualize the bone tissue as histological landmark, and signal in the 642nm channel to visualize the antibody-labeled LYVE-1+ initial lymphatic vasculature network. The lightsheet imaging plane moved from the medial side of the joint towards the lateral side at 1μm intervals to collect a series of 2D images on the sagittal orientation. After mounting the joint onto the tissue adaptor, we utilized tissue autofluorescence in the 488nm channel to define the region of interest for scanning as shown in Fig.2, which resembles a histology slide. We observed the bony tissues, including the distal end of the femur and the proximal end of the tibia and the menisci that composed a knee joint. Within the tibia and femur, the subchondral area with higher signal intensity is sandwiched by lower signal intensity articular cartilage and growth plate (Fig.2A), which we used as the demarcation of the region of interest of the synovial tissues in murine joints. We also observed the patella ligament and the popliteal fossa, which we used as histological landmarks to identify the anterior and posterior of the joints (Fig.2A). We started collecting images when at the separation of the menisci (Fig.2G), and the ending plane when we observed the entirety of the cruciate ligament, which amounts to 550-650 images in the 5-month male wild-type mouse joints, i.e., 550-650μm. We also used the femoral and tibial growth plates to define the proximal and distal boundary of our scan (Fig.2A). Interestingly, while we observed sparse LYVE-1+ signal throughout the joint on 2D images, the distribution of the initial lymphatic network had large variability within a joint, indicating that the quantification of the initial lymphatics is impacted by histological sampling. Thus, examining lymphatic vessel network on 3D images is critical.

**Figure 1.**
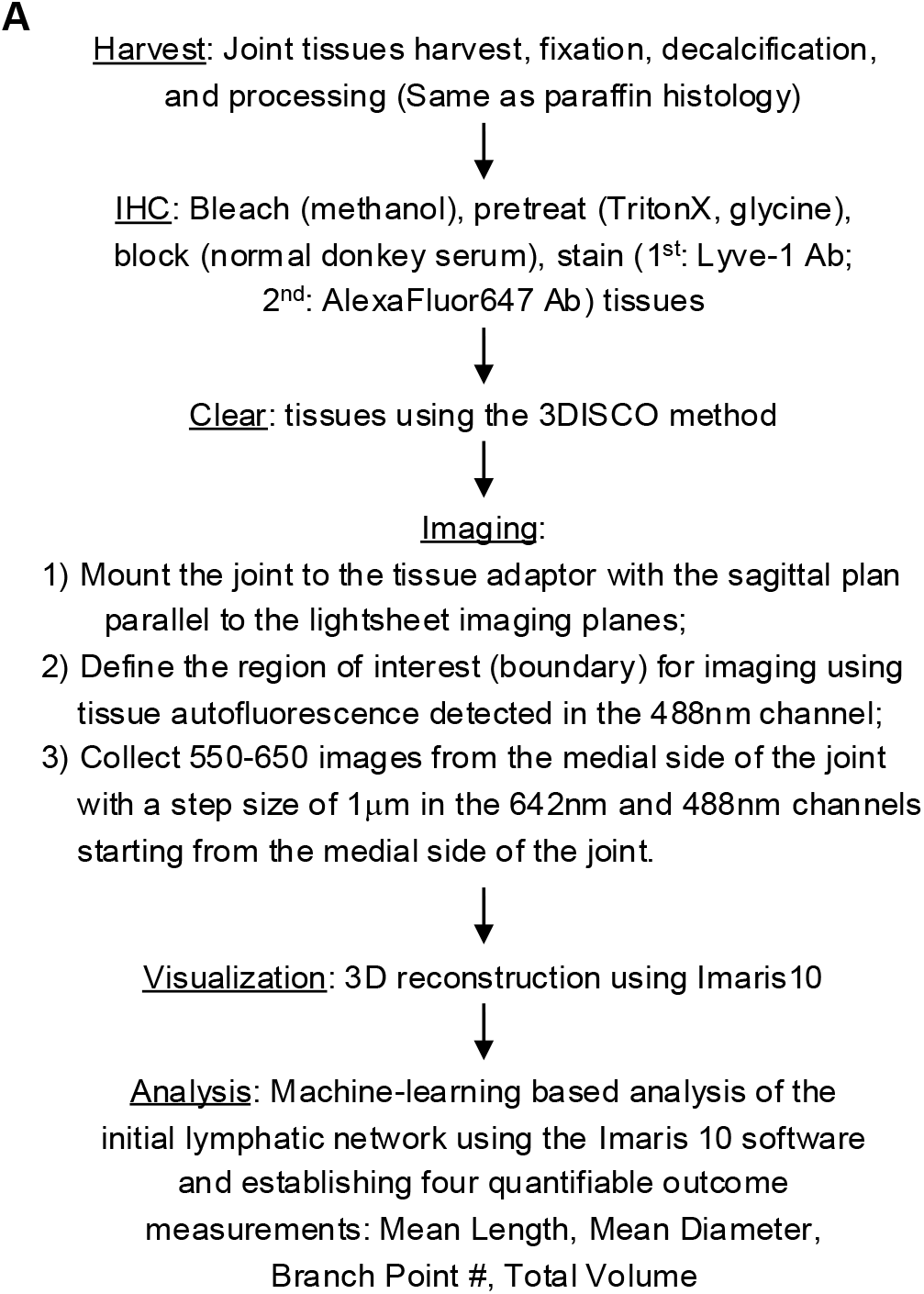
Workflow of lightsheet imaging microscopy for the synovial lymphatic system and quantifiable outcome measurements..

**Figure 2.**
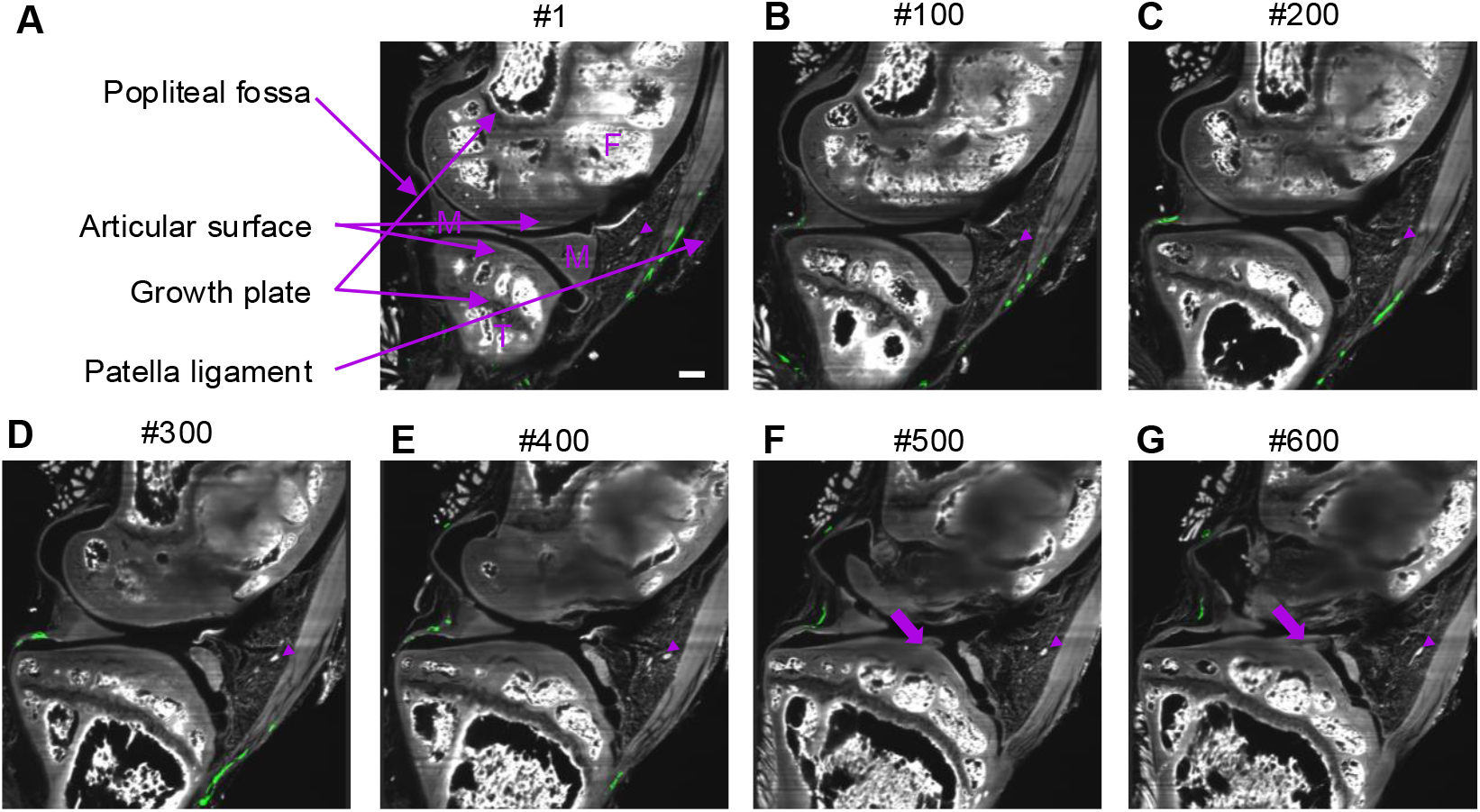
Using tissue autofluorescence to define the region of interest for lightsheet imaging. Knee joints of a 5-m-old C57Bl/6 male mouse were harvested and processed for the lightsheet microscopy of the synovial lymphatic system established in Fig.1. Synovial lymphatic vessels were labeled with LYVE1/AlexaFluor647 antibodies. Joints were mounted in a sagittal position, and a series of 550-650 consecutive images were collected from the medial side of the joint with a step size of 1mm. Fluorescence signal were collected in the AlexaFluor647 channel to visualize the LYVE1-stained lymphatic vessels (green) and GPF channel to visualize the anatomic structure of the joint using the GPF autofluorescence of the bone. (A) Indicated number of the cross sections from the ~600 images collected from the lightsheet microscopy, showing the histology landmarks of a knee joint including the distal end of femur and the proximal end of tibia with articular cartilage, subchondral area and the growth plate, menisci, patella ligament, and popliteal fossa. Histological landmarks are used to define the region of interest for imaging. F=femur, T=tibia, M=meniscus, Scale bar = 200 μm. (A-G) Arrowhead = A blood vessel that is running vertical to 2D imaging plane seen through out the imaging stack (F-G) Arrow= Cruciate ligament attached to the tibial surface is seen in the deeper planes of the images.

### 3D reconstruction showing the initial lymphatic network in the synovial tissue

We thresholded the fluorescence intensity of both 488nm and 642nm channels and performed 3D reconstruction. The 488nm channel revealed the structure of a knee joint (Fig.3A). 3D reconstruction of the 642nm channel LYVE-1+ signal revealed initial lymphatic network on both the anterior and posterior synovial tissues recognized by the patella ligament and popliteal fossa, respectively (Fig.3A). To better visualize the initial lymphatic network, we rotated the 3D reconstructed structure anteriorly (Fig.3B) and posteriorly (Fig.3C). Strikingly, we could observe intricate branching of the initial lymphatic network that is obscured by the sagittal view, highlighting the importance of using whole tissue imaging to visualize the initial lymphatic network. To develop quantifiable outcome measurements, we utilized a machine learning based algorithm in Imaris10 software to identify branching vasculature based on the local intensity contrast of the fluorescence signal of the LYVE-1 staining, which overlaps with the LYVE-1 fluorescence signal (Fig.3D-F), suggesting that the algorithm can successfully predict the vasculature branching. To ensure the accurate modeling of the initial lymphatic network, we used a dynamic range to set the minimum and maximum range of the lymphatic vessels in the training set (Supplementary Fig.1A). Notably, the maximum diameter range for the sham joints is 45.63 ± 2.29 μm, similar to 43.60 ± 3.99 μm in PTOA (*p* = 0.49) (Supplementary Fig.1C). The minimum diameter range for the sham joint is 24.73 ± 4.45 vs significantly smaller 6.40 ± 0.75 μm in PTOA joints (*p* = 0.0021) (Supplementary Fig. 1B). In addition, the branching points were also identified (Fig.3D-F). Thus, it is possible to describe the changes in the joint initial lymphatic network by quantifying the number of branching points/branches, the length and diameter of individual branches, and the total volume of the initial lymphatic network.

**Figure 3.**
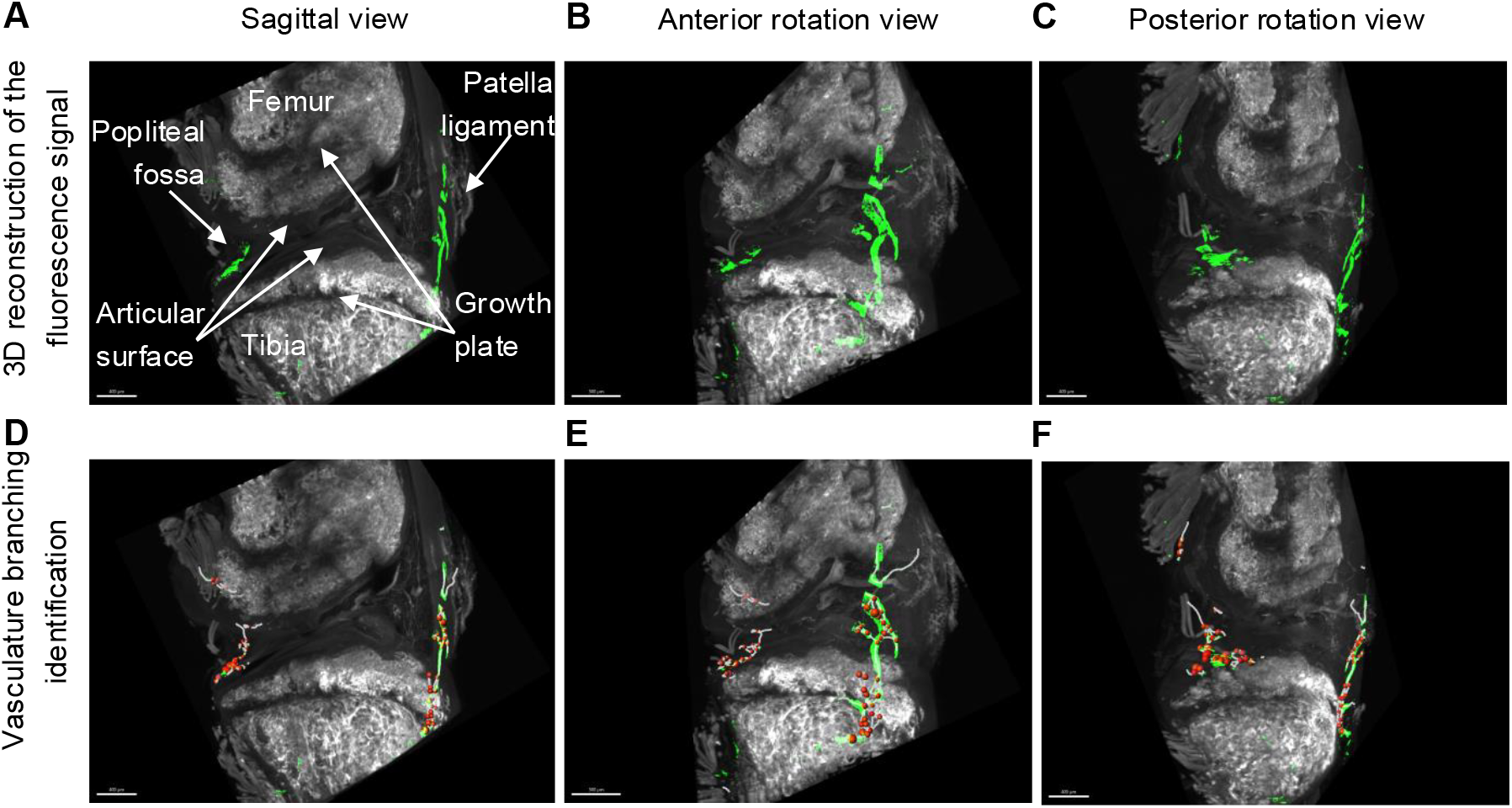
3D reconstruction showing the initial lymphatic network in the synovial tissue. (A-C) Using the fluorescence intensity threshold, we performed 3D reconstruction of the joint tissues in the GFP channel and initial lymphatic network in the 642nm channel. (D-F) Using the machine learning based “filament” algorithm in Imaris10, we performed lymphatic branching analysis and developed quantifiable parameters. Briefly, filaments (the white lines) are identified based on the local intensity contrast of the fluorescence signal of the LYVE1 staining, which overlaps with the LYVE1 fluorescence signal. Branching points are also identified (red dots). The number of branching points/branches, the length of individual branches, and the mean diameter of the individual branches can be used as quantifiable outcome measures to assess the histomorphometry of the synovial lymphatic system. Scale bar = 400 μm.

### The initial lymphatic network of knee joints with post-traumatic OA is visibly different from healthy joints

To validate the lightsheet imaging for synovial lymphatics, we incorporated a group of PTOA joints. We reported previously that PTOA joints have decreased SLS function due to pathological lymphangiogenesis and increased number of lymphatic capillaries revealed by Immunoflourescence/whole slide imaging (IF/WSI) [13, 17]. Consistent with our previous findings, the number of lymphatic vessels is increased in PTOA joints, but the vessels are shorter with lower signal intensity (Fig.4A). After fluorescence intensity thresholding based 3D reconstruction and machine-learning based branch point recognition, we found drastic visible difference between sham and PTOA joint (Fig.4A). Specifically, the lymphatic vessels in PTOA joint are smaller in diameter. Furthermore, when we enlarged and rotated the branching structure of the initial lymphatic, PTOA joint had increased number of branch points, indicated by the red dots (Fig.4B), as well as shorter vessels in length as a consequence (Fig.4B).

**Figure 4.**
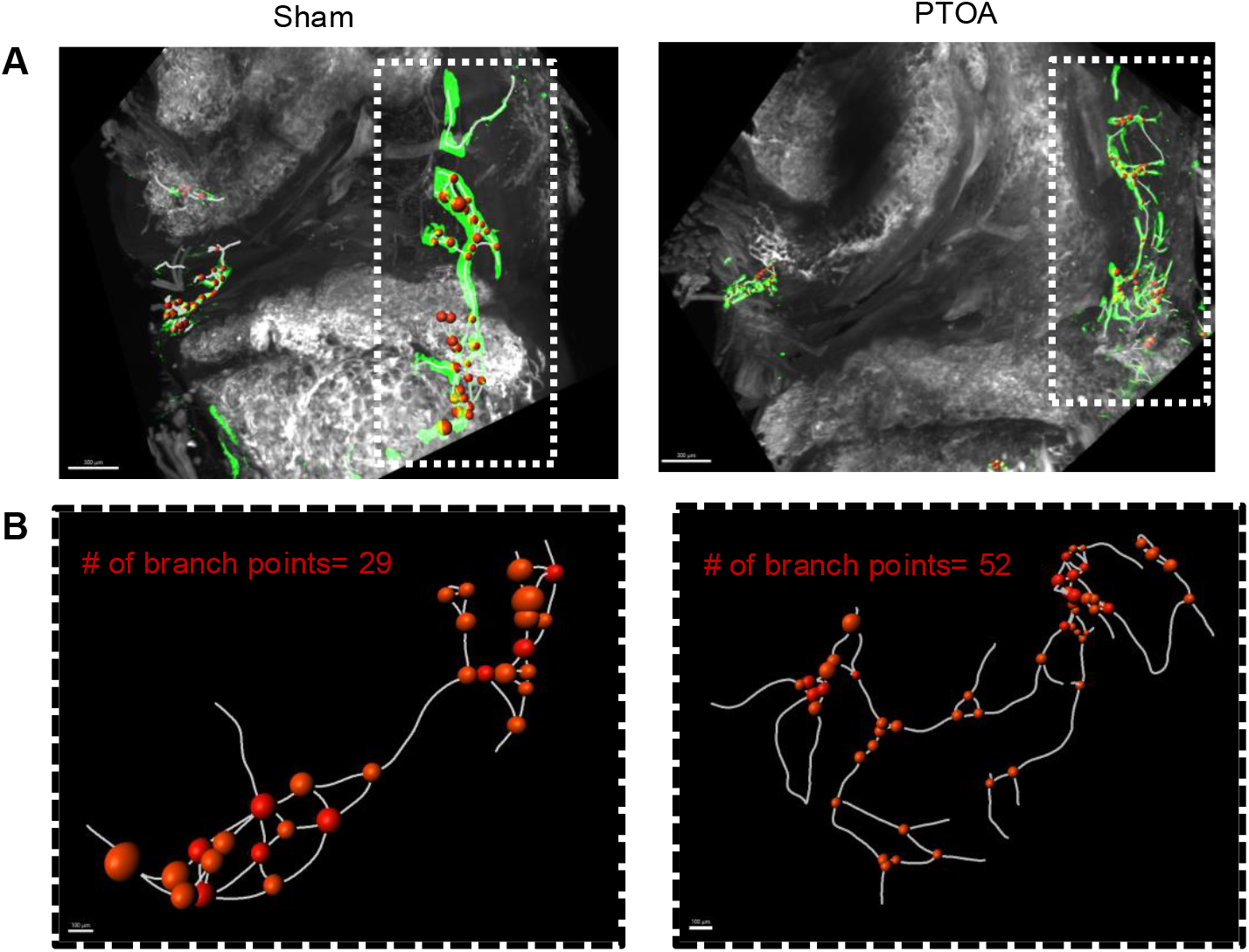
PTOA joints have visibly different initial synovial lymphatic network. (A) 3D reconstruction of the LYVE1 staining showing the lymphatic vasculature (green) in the joint using the “filament” function in Imaris11 (Oxford), which also identifies branch points (red). Noted that the lymphatic vessels in the PTOA joint is thinner but has more branches. (B) Enlarged lymphatic vascular structure in the frontal side of the joint. The structure is rotated for to optimize the visualization the branch points (red balls). The lymphatic vasculature in the frontal section of the joint has 52 branch points in PTOA vs. 29 in sham. The size of the branch points represent the diameter of the lymphatic vessels at the location of branching.

### Lymphatic vessel branch number, length and number as outcome measurement to evaluate the initial lymphatic network

To develop quantifiable outcome measurement based on branch point analysis, we first look at data distribution of the length and diameter of individual branches recognized by the branching model from one sham control and one PTOA mouse. The PTOA joint had 262 individual branches compared to 51 in the control joint (Fig.5). The average length (52.31± 49.94 μm comparted to 143.99 ± 159.88 μm in sham, *p*=0.00026) and diameter (13.16 ± 3.59 μm vs. 29.26 ± 6.25 μm in sham, *p*<0.0001) of the individual branches in the PTOA joint are significantly smaller than control (Fig.5)..

**Figure 5.**
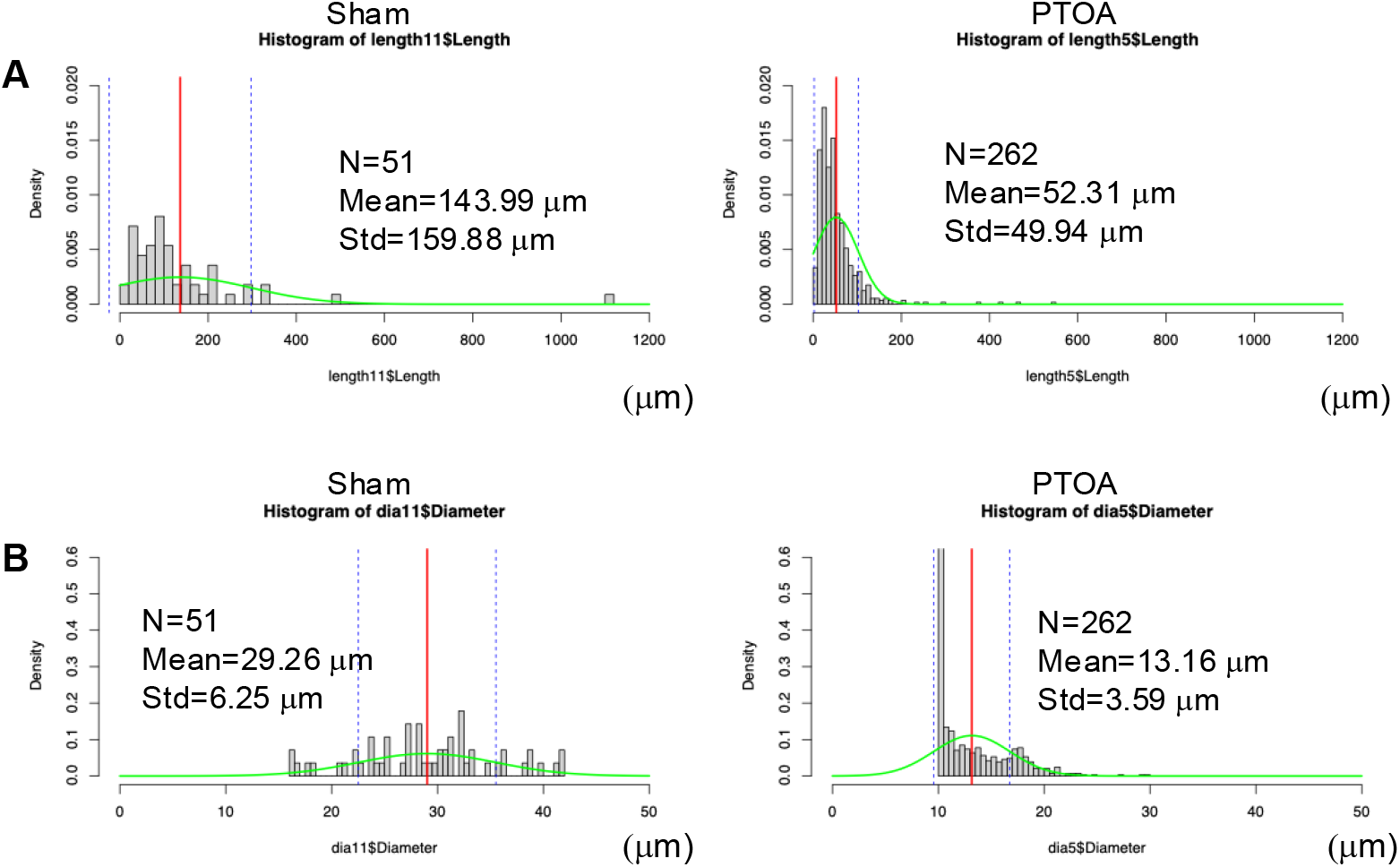
Initial lymphatic network vessels in PTOA joints are shorter in length and smaller in diameter compared to sham joints. (A) Histogram showing the length of individual lymphatic vessel branches. The PTOA joint has more branches with significant shorter branches with 52.31± 49.94 μm comparted to 143.99 ± 159.88 μm in sham, *p* = 0.00026. (B) Histogram showing the average diameter of individual lymphatic vessel branches. Lymphatic vessel branches in the PTOA joint has significantly smaller diameter 13.16±3.59 μm vs. 29.26±6.25 μm in sham, *p*<0.0001. Since the data distribution of the length diameter from one mouse is not normally distributed, the Mann Whitney U test, the non-parametric equivalent of the student t-test, was used analyze the statistical difference between the length/diameter of individual branches in one sham mouse and one PTOA mouse.

**Figure 6.**
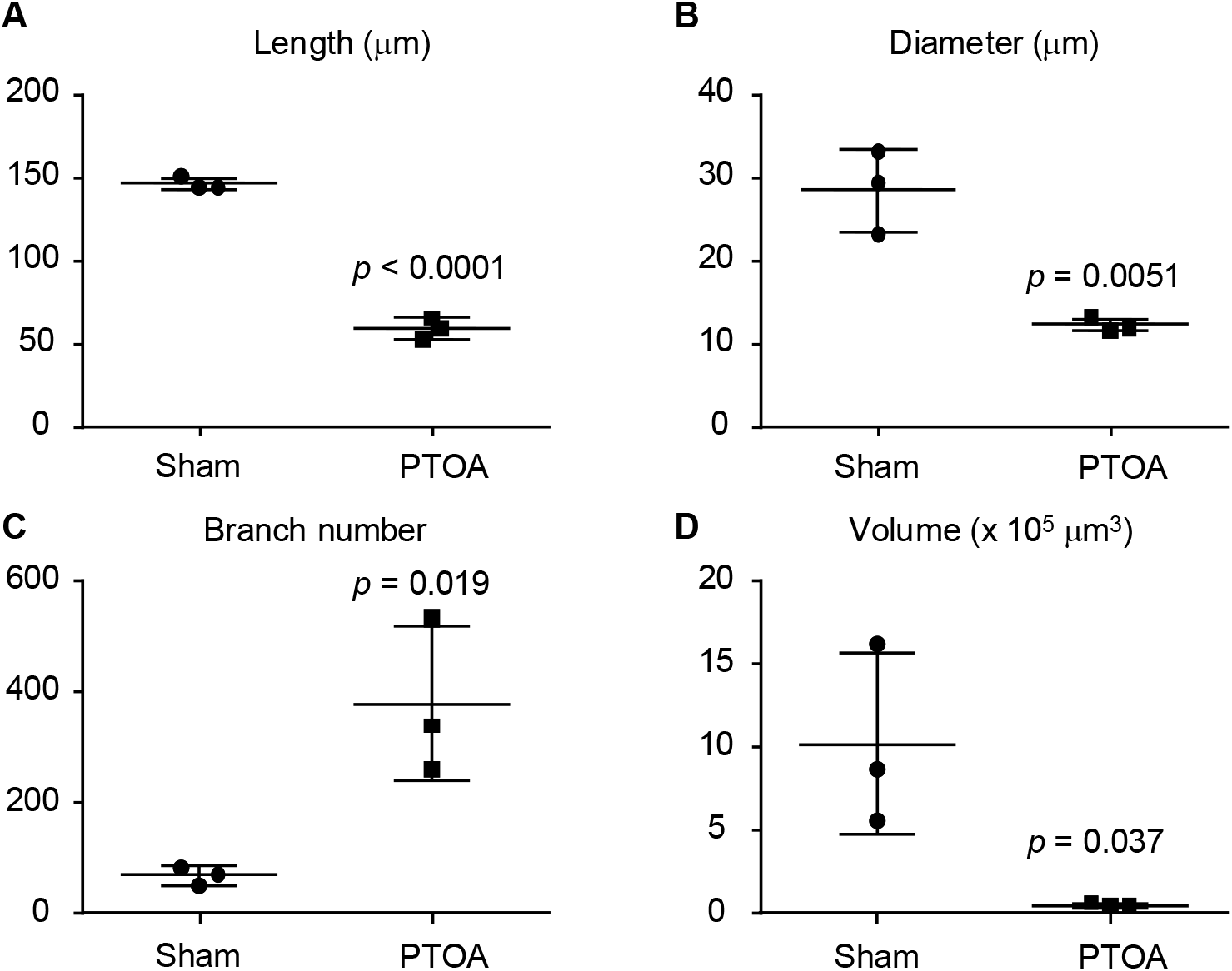
Using four quantifiable parameters to assess the difference in the initial lymphatic network vessels between PTOA and sham control joints. (A) Length: The mean length of all individual branches recognized by Amira in one joint. (B) Diameter: The mean diameter of all individual branches recognized by Amira in one joint. (C) Branch number: The total number of branch points in one joint. (D) Volume: The total volume of the initial lymphatic network of one joint. Student t-test were used to assess the statistical difference between sham and PTOA group. Since the data distribution of the length diameter from one mouse is not normally distributed, the Mann Whitney U test, the non-parametric equivalent of the student t-test, was used analyze the statistical difference between the length/diameter of individual branches in one sham mouse and one PTOA mouse.

### Initial lymphatic network of PTOA joints have shorter branches, branches with smaller diameter, increased branching points, and smaller total volume

To develop quantifiable outcome measurement to assess the initial lymphatic network, we increased the sample size to three joints each group based on power analysis. To streamline the comparison, we took the mean value of the length and diameter individual branches, and found initial lymphatic branches are shorter in length (58.91± 6.50 μm comparted to 146.29 ± 2.48 μm in sham, *p*<0.0001) and smaller in diameter length (12.25± 0.80 μm comparted to 28.51 ± 4.99 μm in sham, *p*=0.0051) compared to control joints. In addition, PTOA joints have increased branching points (378± 239 μm comparted to 69 ± 17 μm in sham, *p*=0.020) but a smaller total volume length (51246.99± 14238.60 μm^3^ comparted to 1019289 ± 544458.32 μm^3^ in sham, *p*=0.037) of the initial lymphatic network. This data is consistent with our previous report that the number of smaller capillary LVs is increased while the larger mature LVS is decreased in early stage PTOA[13], validating the feasibility of the outcome measurements we developed from the SLS lightsheet imaging

## Discussion

In the present study, we established a working protocol for the lightsheet imaging and outcome measurements of the initial lymphatic network in the mouse knee joint. We validated our novel imaging module using samples from the PTOA mouse model.

Over the past decade, our team and others has demonstrated the important role of the SLS in inflammatory arthritis, post-traumatic osteoarthritis, and age-related osteoarthritis (refs). The SLS dysfunction, primarily manifested as decreased clearance of macromolecules, exacerbates arthritis, which is rescued by promoting the SLS function. However, an area of research that requires more attention is the link between SLS function and lymphatic vessel morphology, which is hindered by the plasticity and heterogeneity of the lymphatic system. Cytokines such as TNF, IL1, VEGFC induced by the NFkB pathway during inflammation stimulate lymphangiogensis in inflammatory condition, resulting in increased lymphatic capillary but not mature lymphatic vessels, which is validated [13, 17] by the increased branching and decreased diameter in our study. We have also shown the decrease of SLS function in natural aging joint associated with decreased VEGFC level, which is an independent pathway from the inflammatory arthritis. Specifically, we observed decreased SLS function via IVIS-Dextran imaging, including the accumulation of macromolecules in the joint space, decreased macromolecules in the initial lymphatic network and the upstream DLN. Counterintuitively, we have not observed morphological change such as decreased number of lymphatic vessels in the joint. Using the lightsheet imaging of the initial lymphatic network could reveal otherwise undetected changes in the SLS system.

In the present study, we developed three parameters to evaluate the initial lymphatic network of the SLS, including the number of branches, the average length, and diameter of the lymphatic branches. We combined the analysis of the anterior portion (the patella ligament side), and the posterior (the popliteal fossa side) of the synovium. LYVE1+ staining is higher in the anterior portion (Fig.2A-D) compared to the posterior portion (Fig.2E-F) in the more superficial levels of the joint (larger menisci) than in the deeper levels (cruciate ligaments). A more comprehensive investigation of location-specific initial lymphatic distribution and morphology may help unravel the mechanisms underlying SLS related pathology, especially in conditions such as natural aging. In addition, we cut off our analysis at the two growth plates of the femur and tibia, according to the standard operation protocol of histomorphometry analysis of the synovial tissues. But we did observe the initial lymphatic vessels extended beyond the boundary of our analysis (Fig.2C&D). It may be interesting to probe the initial lymphatic network to more proximal and distal region of the joint.

Besides the LYVE1+ staining and the histological landmarks, we can also observe other structures in the 488nm autofluorescence channel, including the fat pad and vessel-like structures within the fat pad (Fig.2, arrowhead), which is part of the sublining synovial tissues. While articular cartilage is avascular, the synovial tissue receives vascular supply by a highly anastomoses system, termed the periarticular plexus derived from multiple arteries and veins[18]. In a healthy human, the periarticular plexus is distributed with 90 μm deep of the lining synovial tissue, and is at extended deeper into the sublining synovial tissue at 165 μm deep [18]. Interestingly, in a study in 2013 where the authors used contrast enhanced CT to investigate the vasculature system of the hindlimb of wildtype mice, the anterior part of the mouse knee joint has a lack of major arteries [19]. Thus, with the power of lightsheet microscopy, it will be interesting to probe the correlation between lymphatic vasculature and blood vasculature and potential crosstalk to generate a comprehensive atlas in the future.

In the present study, we focused on the initial lymphatic network of murine knee joint, where we previously link to arthritis pathogenesis [15, 17, 20]. Our methodology is significant because we might adapt it into other joints such as ankle and hand joints, as well as other musculoskeletal system such as bone and tendon. One technical challenge of applying knee lightsheet to other joints or musculoskeletal tissues is the balance between decalcification and antigen preservation. While establishing the protocol, we have tried to shorten our decalcification time from standard 14 days to 3 days to maximize the antigen preservation for immunolabeling. However, we found that high calcium content in the tissue impede the tissue clearing and consequently generate shadows when the lightsheet plane moved across the tissue. Indeed, some part of the subchondral area of the femur was obstructed by the shadow in the 488nm channel (Fig.2D-E) the deeper the lightsheet plane is and due to the thicker bone tissues of the femur compared to the tibia. Thus, optimization is needed if we want to examine the correlation between the subchondral area or other deep tissues. Another potential obstacle of applying knee SLS lightsheet to other organ is the depth of the lymphatic vessel distribution. The initial lymphatic network is distributed within the synovium. However, previous studies have shown that the collecting lymphatic vessels of the ankle, which are two major lymphatic vessels parallel to a vein, run superficially on top of the muscle tissues and drain into the popliteal lymph node in the popliteal fossa. A general principle for tissue harvest for the lightsheet protocol is to remove all the tissues that are not region of interest. In our protocol we carefully removed as much as possible, because muscle tissue would result in high non-specific binding during immunostaining. If we want to generate a complete atlas of the SLS, which includes the initial lymphatic network, the collecting lymphatic vessel, and the draining lymph node, we will need to optimize the tissue dissection process.

There are several limitations in our study. 1) We only imaged the initial lymphatic network in half the joint: from the medial side to the cruciate the ligament based on our standard operation protocol of mouse knee histology (ref). In the future, we should also scan the knee from the lateral side to generate a comprehensive atlas for whole knee. 2) We stained the initial lymphatic network with anti-LYVE-1 antibody. We should optimize our protocol using other antibodies that specifically recognize lymphatic endothelial cells such as Prospero homeobox protein 1 and Podoplanin. 3) Anti-LYVE-1 antibody stains all lymphatic vessles. We will need to include anti-smooth muscle actin antibody to distinguish lymphatic capillary and mature lymphatic vessels.

## Conclusion

Lightsheet imaging of murine knee joints is a powerful and promising tool to study the synovial lymphatic system and arthritis. Initial lymphatic network of the PTOA joint has increased branching with smaller and shorter branches.

## Data Availability Statement

The raw data supporting the conclusions of this article will be made available by the authors, without undue reservation.

## Author Contributions

LinX, HZ and LX designed the work. LinX performed experiments and analyses. LinX and LX wrote the original manuscript. All the authors discussed the result and approved the manuscript.

## Acknowledgements

The lightsheet microscope was housed in our facility URMC Center for Advanced Microscopy & Nanoscopy (RRID:SCR_023177)

## Figure Legends

**Supplementary Figure 1. The minimum and maximum lymphatic vessel diameter threshold was set for branching analysis**. To ensure accurate modeling the branching network, the minimum and maximum diameter was set differentially for individual mouse by scrolling through the z-stack using a ruler tool. (A) Bar graph showing the range of the diameter set for branching analysis. (B) The minimum diameter was set significantly lower in the PTOA group compared to the sham group. (C) The maximum diameter was set comparably in the PTOA group compared to sham group. Student t-test was used analyze the statistical difference between the length/diameter of individual branches in one sham mouse and one PTOA mouse

