## Supplementary Figure 1 for "Using lightsheet microscopy to investigate the initial lymphatic network in the murine knee joints"

### Slide 1
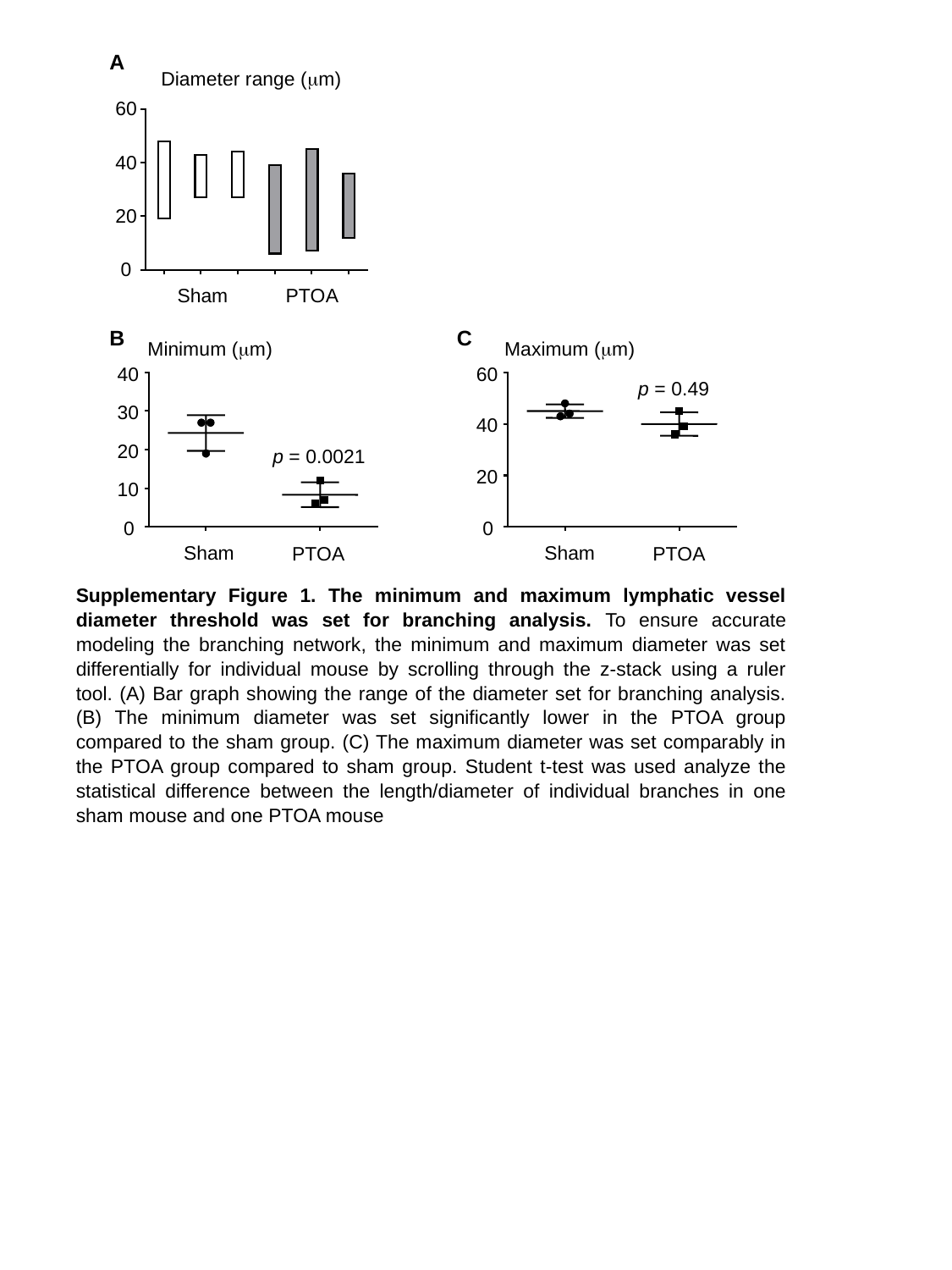

A
Diameter range (μm)
60
40
20
0
Sham
PTOA
B
C
Minimum (μm)
Maximum (μm)
40
60
p = 0.49
30
40
20
p = 0.0021
20
10
0
0
Sham
Sham
PTOA
PTOA
Supplementary Figure 1. The minimum and maximum lymphatic vessel diameter threshold was set for branching analysis. To ensure accurate modeling the branching network, the minimum and maximum diameter was set differentially for individual mouse by scrolling through the z-stack using a ruler tool. (A) Bar graph showing the range of the diameter set for branching analysis. (B) The minimum diameter was set significantly lower in the PTOA group compared to the sham group. (C) The maximum diameter was set comparably in the PTOA group compared to sham group. Student t-test was used analyze the statistical difference between the length/diameter of individual branches in one sham mouse and one PTOA mouse
